# Functional Connectivity Differences Underly Variable Centralized Symptom Relief After Hysterectomy

**DOI:** 10.64898/2026.09.26.754679

**Authors:** Natalie McLain, Chelsea Kaplan, Steven Harte, Melissa Lenert, Julia Evanski, Esmeralda Hidalgo-Lopez, Maximillian Egan, Eric Ichesco, Richard E. Harris, Jason Kutch, Daniel Clauw, Sawsan As-Sanie, Andrew Schrepf

## Abstract

Approximately 1 in 4 women with chronic pelvic pain (CPP) report persistent pelvic pain after hysterectomy. Centralized symptoms—fatigue, multisite pain, and sleep disturbance, quantified using the Fibromyalgia Survey Score (FSS)—often co-occur with this risk, but the trajectory of centralized symptom burden after surgery, independent of pelvic pain outcome, is not well understood. Whether presurgical differences in central nervous system pain processing distinguish women whose centralized symptoms improve (improvers) from those whose do not (non-improvers) has not been tested. In this observational study, we assessed whether individuals who do and do not report improvement in FSS scores six months post-operatively differ in their baseline/pre-operative phenotypes and brain function. Sensory sensitivity scores were significantly higher in non-improvers than in improvers at baseline. We identified no significant, presurgical differences in age, average pelvic pain intensity, widespread pain, or pain interference between improvers and non-improvers. Functional connectivity was examined at rest and during the application of a distal, sustained pressure pain. We identified no significant group differences at rest. During sustained pressure pain, non-improvers showed greater connectivity than improvers between primary somatosensory cortex and posterior thalamic seeds with frontal, motor, cerebellar and limbic regions, but lower connectivity between the periaqueductal gray and subgenual cingulate cortex. Non-improvers also showed a greater pain-evoked increase in connectivity between the pelvic and leg representations of the primary sensory cortex and other sensorimotor regions in the pain-versus-rest contrast. Overall, non-improvers showed increased functional connectivity between regions associated with the multisensory integration of ascending pain, and decreased connectivity between regions associated with descending pain modulation. In exploratory analyses, some of these connections were additionally associated with symptom burden: sensory sensitivity, number of painful regions, and pain interference. This work may inform mechanisms of persistent symptom burden and guide surgical decision-making in patients with high central sensitization.

## Introduction

Chronic pelvic pain (CPP) is a debilitating condition affecting an estimated 15-26% of women globally,^1^ and hysterectomy is a commonly pursued treatment for cases unresponsive to conservative therapy.^2–4^ Most women report significant pelvic pain relief, but a subset (12-68%) report persistent pain following surgery.^5^ Nociplastic pain—pain arising from altered central nervous system processing—is an established mechanism underlying CPP^2^ and is associated with a higher risk of persistent pelvic pain after hysterectomy.^6,7^ Comorbid symptoms of nociplastic pain, including fatigue, poor sleep, and cognitive impairment, are common among women with CPP and meaningfully impair quality of life. Thus, nociplastic symptom burden is relevant not only to the likelihood of pelvic pain relief after surgery, but also to patients’ overall quality of life and global perception of postoperative improvement. However, little is known about how these symptoms themselves respond to hysterectomy or whether presurgical differences in central nervous system function distinguish women whose nociplastic symptoms improve from those whose symptoms persist.

Rather than a single disease category, nociplastic pain is increasingly understood as a dimensional phenotype shaped by altered sensory processing, large-scale brain-network dysfunction, and abnormal engagement of descending pain modulatory systems.^8^ Current models of nociplastic pain emphasize that similar symptom profiles can exist with or without ongoing peripheral pathology.^8^ This distinction is particularly relevant to hysterectomy, which is a peripherally directed intervention intended to remove a putative source of nociceptive input. Although greater nociplastic pain symptom burden is associated with poorer pain outcomes, women with substantial nociplastic symptoms may nevertheless experience valued degrees of pain relief following hysterectomy.^6^ Together, these findings point to clinically important heterogeneity in the extent to which a patient’s symptom burden is maintained by ongoing peripheral nociceptive input versus more centrally sustained processes.

We previously showed that roughly two-thirds of people undergoing total hip or knee arthroplasty experience considerable improvement in pain across multiple body regions, fatigue, dysregulated sleep, and cognitive fog, six months after the procedure, whereas roughly one third experience no such benefit — despite high rates of symptomatic joint pain improvement in both groups.^9^ We termed these groups “bottom-up” and “top-down” forms of nociplastic pain, respectively, given the distinctly different trajectories following removal of nociceptive input. We hypothesized that non-painful sensory sensitivity may help distinguish between nociplastic subtypes. But as we had no measures of these symptoms, we were unable to identify robust differences between groups that would allow us to distinguish nociplastic subtypes prior to surgery.

Using an expanded phenotyping battery using neuroimaging and sensory sensitivity, we sought to identify neural and symptomatic correlates differences in nociplastic subtypes. Neuroimaging offers a way to measure differences in central nervous system activity within the proposed bottom-up and top-down nociplastic pain framework. Across nociplastic and chronic overlapping pain conditions, altered activity has been reported in somatosensory cortex,^11-13^ thalamus,^14-16^ and periaqueductal gray (PAG),^16,17^ as well as in broader interactions among default mode, salience, and sensorimotor systems.^8^ Presurgical differences within these systems may therefore distinguish patients whose nociplastic-associated symptoms remain centrally sustained after surgery from those whose symptoms improve when a peripheral nociceptive source is removed.

The present study examined females scheduled to undergo hysterectomy for CPP at a tertiary pain clinic. Our primary aim was to determine whether presurgical functional connectivity from *a priori*, pain-relevant brain regions, assessed during rest and sustained pressure pain functional magnetic resonance imaging (fMRI), distinguished patients whose nociplastic-associated symptoms improved after surgery from non-improvers. We hypothesized that non-improvers would show altered engagement of ascending sensory and descending modulatory circuitry, including S1, thalamus, and PAG, consistent with nociplastic symptoms maintained at supraspinal levels despite removal of peripheral nociceptive input.

## Materials and methods

### PARTICIPANTS

Participants were drawn from a prospective observational study: Peripheral and Central Nervous System Mechanisms of Persistent Post-Hysterectomy Pain. This study enrolled 423 females scheduled to undergo hysterectomy for benign indications at the University of Michigan between December 2017 and July 2022. Baseline study procedures were completed within 30 days before hysterectomy, and follow-up questionnaires were collected after surgery, including at 6 months. The present analysis included only the subset of participants with CPP who completed preoperative functional MRI and had available 6-month symptom data.

For the parent study, participants were eligible if they were 21 to 70 years of age, scheduled for hysterectomy for a benign, non-cancer indication, and able to read and speak English to complete informed consent and patient-reported outcome (PRO) measures. Participants were excluded for severe medical or psychiatric conditions, concurrent major surgery (e.g. bowel resection) or other circumstances judged by the study team to make participation unsafe or interfere with interpretation of study results. Participants were additionally excluded for concurrent participation in therapeutic trials, pregnancy and/or lactation, illegal drug use or substance abuse within the past two years.

The subset of participants in this analysis were part of the chronic pelvic pain subset of the parent study. To be assigned to this group, participants were required to report pelvic pain for at least 6 months, pelvic pain on at least 5 days per month, average pelvic pain in the prior month of at least 4 on a 0–10 numerical rating scale (NRS), and pain localized to the anatomic pelvis that was not limited to dyspareunia or dyschezia. Participants with suspected or previously diagnosed endometriosis were eligible for the parent chronic pelvic pain/endometriosis cohort as well as the subset analyzed here.

Participants in this analysis additionally were required to meet inclusion/exclusion criteria for neuroimaging and quantitative sensory testing (QST), including right-hand dominance, no contraindications to MRI or pressure-cuff testing, able to lie supine and still for the MRI session, and normal or corrected to normal vision for viewing scanner instructions. Participants were asked to refrain from over-the-counter pain medications, alcohol, and nicotine on the day of MRI and QST, and from exercise likely to cause muscle or joint soreness for the 48 hours before their visit. Participants using adjunctive pain medications with potential effects on fMRI or QST were studied off these medications when feasible or were asked to remain on a stable dose before testing.

All participants provided written informed consent approved by the University of Michigan Institutional Review Board [HUM00117473].

### PHENOTYPING BATTERY

Patients completed a battery of PRO measures in the 30 days prior to surgery and 6 months post-surgery. Measures in the current analysis are described below.

#### Pelvic pain intensity

The following item was used to assess pelvic pain intensity: “In the past 3 months, how would you rate your AVERAGE pelvic pain, cramping, or discomfort you have experienced? (0-10).”

#### Pain interference

The PROMIS pain interference short form 8a was used to assess pain interference (0-40).^18^

#### Nociplastic pain

The American College of Rheumatology 2011 Fibromyalgia Survey Score (FSS) was used to measure nociplastic-associated pain and symptoms.^19^ The FSC comprise two components: the Widespread Pain Index (WPI), which assesses the number of painful body regions (0-19), and the Symptom Severity score, which assesses fatigue, trouble thinking, and sleep disturbance (0-12). The total FSC score ranges from 0 to 31, with higher scores indicating greater nociplastic symptom burden. Although the FSC was originally developed for epidemiologic classification of fibromyalgia, it has also been used dimensionally as a proxy measure of nociplastic pain features across chronic pain conditions^6,20^ given its association with specific symptom profiles^5,20-22^

#### Spatial distribution of pain

The WPI subscale from the FSC^19^ (see above) was used separately from the FSC score to assess number of painful body regions. The map is made up of 19 sites: right and left jaw, shoulder, upper arm, lower arm, hip/buttock/trochantor, upper leg, and lower leg, as well as single regions for neck, upper back, lower back, chest and abdomen.

#### Sensory sensitivity

Self-reported sensory sensitivity was assessed using the Complex Medical Symptom Inventory (CMSI). This survey is composed of 41 items asking about the presence of functional symptoms for 3 months out of the past year.^23^ In this study, we used the subset of items that capture sensory sensitivity to nonpainful environmental stimuli (CMSI35-Sensitivity to certain chemicals, such as perfumes, laundry detergents, gasoline and others; CMSI36-Sensitivity to sound; CMSI37-Sensitivity to odors; CMSI39-Frequent sensitivity to bright lights), with scores ranging from 0 (no sensitivities)-4 (heightened sensitivity to all four sensory modalities).^24,25^

### SAMPLE SELECTION

From the original dataset of 423 females, 72 completed the 6-month follow-up visit and had complete baseline neuroimaging data. These patients were divided into two groups based on the change in their FSC scores from baseline to 6 months post-hysterectomy. Participants with greater than 50% reduction in FSC were classified as improvers, whereas participants with less than 50% reduction were classified as non-improvers. Participants were then matched across improver and non-improver groups within two points of their baseline FSC score using the “optmatch” package in R for a final sample of 54 participants, with 27 improvers and 27 non-improvers. A subset of 42 participants, with 21 improvers and 21 non-improvers, had both resting state and pressure pain fMRI data.

Matching improvers and non-improvers on baseline FSC, rather than adjusting for the score as a covariate, provides distribution-level control that does not depend on correctly specifying the relationship between FSC and functional connectivity. Because FSC is arithmetically coupled to the group definition (percent change from baseline) and subject to regression to the mean, baseline scores would be expected to differ between groups; matching balances this variable across groups without assuming a linear covariate effect, supporting the interpretation that observed connectivity differences reflect improver status rather than baseline symptom burden.

### MRI PROTOCOL

#### MRI Acquisition

Neuroimaging was performed before hysterectomy with participants positioned supine and entering the scanner headfirst. Images were acquired using a General Electric MR750 Discovery 3.0T scanner (Modular Unit MRI Machine) and a General Electric 3.0T Ultra-High Performance (UHP3T) scanner, both with a Nova Medical 32 channel head coil. Participants were asked to keep their eyes open and fixed on a screen with a white plus sign on a dark gray background.

A high-resolution T1-weighted anatomical image was acquired for functional-image registration and anatomical normalization using a sagittal three-dimensional magnetization-prepared rapid gradient-echo sequence. Acquisition parameters included an inversion time of 1060 ms, flip angle of 8°, field of view of 256 × 256 mm, acquisition matrix of 256 × 256, 208 contiguous sagittal slices, and 1-mm isotropic spatial resolution. Repetition time (TR) was 2500ms and echo time (TE) was 2ms.

Functional images were acquired using a multiband gradient-echo echo-planar imaging sequence with repetition time/echo time = 800/30 ms, flip angle = 52°, field of view = 216 × 216 mm, acquisition matrix = 90 × 90, 60 contiguous axial slices, slice thickness = 2.4 mm, no interslice gap, and multiband acceleration factor = 6, yielding 2.4-mm isotropic voxels. Slices were acquired in interleaved order.

Two resting state fMRI scans were included in this analysis, each approximately 6 minutes in duration. During the first scan, participants underwent resting-state imaging with no experimental stimulus. The second scan, acquired approximately 50 minutes later in the same fMRI session, was conducted during application of tonic pressure pain. For this condition, an MRI-compatible pneumatic cuff was placed around the left gastrocnemius muscle. Before scanning, cuff pressure was calibrated individually to each participant’s Pain40 threshold, defined as the pressure intensity that evoked a pain rating of ∼40/100 on a NRS. The calibrated pressure was then applied continuously during this scan.

#### fMRI and T1 preprocessing

Functional data preprocessing conducted through fMRIPrep included co-registration to structural T1 (bbregister) and realignment (mcflirt, FSL 5.0.9). A deformation field to correct for susceptibility distortions was estimated based on SDCFlows’ fieldmap-less approach.^26–28^ Data was additionally normalized to MNI standard space (ANTs 2.2.0), and resampled to 2mm isometric voxels. No slice-timing correction was performed.

The preprocessed fMRIPrep output were entered into the CONN Toolbox: CONN^29^ (RRID:SCR_009550) release 22.a^30^ and SPM^31^ (RRID:SCR_007037) release 12.7771. The first 10 scans in each functional run were removed. Functional data were coregistered to a reference image (first scan of the first session) using a least squares approach and a 6 parameter (rigid body) transformation without resampling.^32^ Functional data were smoothed using spatial convolution with a Gaussian kernel of 6 mm full width half maximum (FWHM). Last, potential outlier scans were identified using ART^33^ as acquisitions with framewise displacement above 0.9 mm or global BOLD signal changes above 5 standard deviations,^34,35^ and a reference BOLD image was computed for each subject by averaging all scans excluding outliers.

Denoising was performed simultaneously and included the following steps: regression of potential confounding effects characterized by white matter timeseries (5 CompCor noise components), CSF timeseries (5 CompCor noise components), motion parameters and their first order derivatives (12 factors),^36^ outlier scans (below 18 factors),^34^ session and task effects and their first order derivatives (4 factors), and linear trends (2 factors) within each functional run, followed by bandpass frequency filtering of the BOLD timeseries^37^ between 0.008 Hz and 0.09 Hz. CompCor^38,39^ noise components within white matter and CSF were estimated by computing the average BOLD signal as well as the largest principal components orthogonal to the BOLD average, motion parameters, and outlier scans within each subject’s eroded segmentation masks.

#### Region of interest to whole brain connectivity analysis

We selected12 *a priori* regions of interest (ROIs) for ascending and descending pain signals based on past literature in pain imaging. Anterior and posterior bilateral thalamus, a total of four seeds, were defined using the Melbourne Subcortex Atlas, Stage 1 (https://github.com/yetianmed/subcortex).^40^ Six regions in S1 were included in this analysis. The first was identified as important for widespread pain in chronic pelvic pain^10^ and later validated as a region predicting the development of widespread pain in a large, longitudinal pediatric dataset.^11^ Three pelvic representation seeds were derived from a previous paper examining both healthy individuals and those with chronic pelvic pain.^41^ One seed representing the stimulated pressure pain site on the left leg (Table 1, seed 5) was derived from a previous publication on the same paradigm.^12^ A final, S1 seed for the left arm was selected to represent a body region not relevant to the chronic pain condition nor the stimulation paradigm. This was included as a form of specificity check for the other S1 seeds that were relevant to pain experienced by the participants, either through their chronic pain condition and/or the pressure cuff stimulation. All six seeds were defined by placing 5mm radius spheres centered on peak coordinates from the previous studies. Finally, two seeds were used to define the periaqueductal grey (PAG) based on previous literature demonstrating PAG ROI definition strongly effects the results of imaging analyses.^42^ Both the MNI centroid and hand traced ROI from Fenske et al., 2020 were entered into this analysis. See Table 1 for a full list of ROI details including centroid coordinates and sources.

**Table 1.** Seeds tested in the seed-to-whole brain analysis. Includes a seed number to clarify ambiguous/overlapping seed names in-text.

| Seed # | Seed label | type | centroid |  |  | source |
| --- | --- | --- | --- | --- | --- | --- |
|  |  |  | x | y | z |  |
| 1 | WidespreadS1-rh | 5mm sphere | 19 | -27 | 71 | Kutch et al. 2017 |
| 2 | S1Pelvic_R1 | 5mm sphere | 18 | -40 | 60 | Bagarinao et al., 2014 |
| 3 | S1Pelvic_R2 | 5mm sphere | 44 | -24 | 56 | Bagarinao et al., 2014 |
| 4 | S1Pelvic_L | 5mm sphere | -26 | -36 | 60 | Bagarinao et al., 2014 |
| 5 | S1LeftLeg | 5mm sphere | 8 | -38 | 68 | Kim et al 2015 |
| 6 | S1LeftArm | 5mm sphere | 29 | -33.4 | 68.8 | Roux et al., 2017 |
| 7 | pTHA-lh | atlas |  |  |  | Tian et al. 2020 |
| 8 | aTHA-lh | atlas |  |  |  | Tian et al. 2020 |
| 9 | pTHA-rh | atlas |  |  |  | Tian et al. 2020 |
| 10 | aTHA-rh | atlas |  |  |  | Tian et al. 2020 |
| 11 | PAG_sphere | 5mm sphere | 4 | -26 | 14 | Fenske et al. 2020 |
| 12 | PAG_trace | hand trace<br>(previously<br>published) |  |  |  | Fenske et al. 2021 |

First-level seed-to-voxel functional connectivity maps were estimated separately for each condition. Functional connectivity was represented by Fisher-transformed bivariate correlation coefficients between the mean BOLD time series from each seed and the time series of every voxel in the brain. For condition-specific estimates, individual scans were weighted by boxcar regressors representing the experimental condition, convolved with the canonical hemodynamic response function.

### SECOND-LEVEL fMRI ANALYSES

Second-level seed-to-voxel analyses were conducted in CONN using general linear models. Three analyses were performed. First, we tested group differences in connectivity during the rest condition using all participants with available rest data (“rest”; n=54). Second, we tested group differences in connectivity during sustained painful pressure using all participants with available painful pressure data (“pain”; n=42). Third, we tested whether pain-evoked changes in connectivity between rest and pain differed between groups among participants with both rest and pain scans (“pain vs rest”; n=42).

For the rest and pain analyses, condition-specific connectivity maps were entered into separate second-level models comparing improvers and non-improvers. Age and mean framewise displacement were included as covariates of no interest. These models tested adjusted between-group differences in seed-to-voxel connectivity within each condition.

For the pain vs rest analysis, rest and pain connectivity maps were entered together in a repeated-measures second-level model. Group was modeled as the between-subjects factor, condition was modeled as the within-subject factor, and age and mean framewise displacement were included as covariates of no interest. This model tested the group-by-condition interaction, identifying regions where the change in connectivity from rest to sustained pressure pain differed between improvers and non-improvers.

For all three analyses, voxel-level significance was set at p < 0.001, and cluster-level significance was set at p < 0.05, false discovery rate corrected for multiple comparisons. Directionality of significant effects was determined from extracted Fisher-transformed connectivity values.

### POST HOC CLINICAL ASSOCIATION ANALYSIS

Fisher-transformed connectivity values were extracted from significant clusters identified across the rest, pain, and pain vs rest analyses. Extracted values were imported into R, version 4.5.1. Box-and-whisker plots were generated to visualize connectivity values by group. Exploratory Spearman correlations were then used to examine associations between extracted connectivity values and three baseline clinical characteristics: pain interference, number of painful body regions, and sensory sensitivity. Spearman correlation analyses were conducted separately within improvers and non-improvers to characterize whether brain-clinical relationships differed by post-surgical symptom trajectory.

## Results

A total of 54 participants were included in the primary fMRI analysis, including 27 improvers and 27 non-improvers. The only tested variable that showed a significant group difference at baseline was sensory sensitivity, with a mean baseline score of 1.15±1.41 in nonimprovers and 0.56±1.15 in improvers. All other variables did not significantly differ between the groups at baseline: the mean number of painful body sites was 2.48±2.23 in non-improvers and 2.59±2.42 in improvers, mean baseline pelvic pain was 5.70±1.68 for non-improvers and 5.67±2.65 for improvers and mean baseline pain interference was 23.48±8.3 in nonimprovers and 21.59±9.3 in improvers. Change scores showed a mixture of significant and nonsignificant results. Change in FSC from baseline to 6 months after hysterectomy significantly differed between groups, with improvers showing a mean reduction of -6.67±3.54 compared with -0.93±3.68 in non-improvers (p < .001; Figure 1a). There was no significant difference in the pelvic pain intensity change from baseline to follow-up, with an average reduction of -4.89±1.63 in non-improvers and - 5.56±2.56 in improvers. However, improvers did see significantly larger reduction in their widespread pain index, with improvers experiencing an average of -2.04 ±2.31 fewer body sites following surgery and only 0.08±3.26 in non-improvers (*p*=0.044) (Figure 1b). Further information on group characteristics at baseline and follow-up can be found in Table 2.

**Figure 1.**
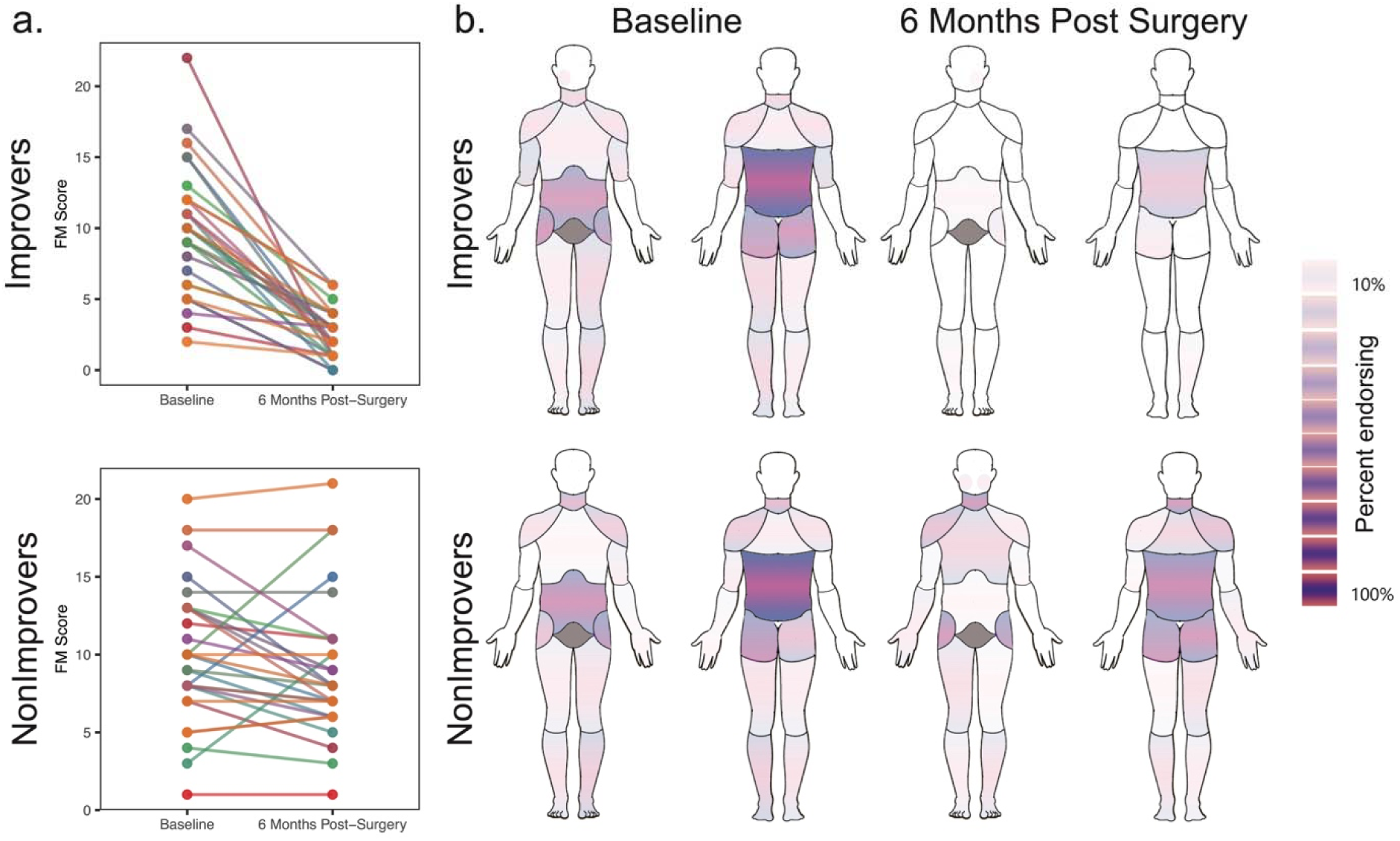
Characterizing improvers and non-improvers. Panel **a** shows individual trajectories of the Fibromyalgia Survey Score (FSS) from baseline to 6 months after hysterectomy for every participant in the improver and non-improver groups. Panel **b** shows the percent of individuals in each group endorsing pain in a regionalized body map before and 6 months following hysterectomy. The map is based on the Widespread Pain Index (WPI), which is made up of 19 sites: Right and left jaw, shoulder, upper arm, lower arm, hip/buttock/trochantor, upper leg, and lower leg, as well as single regions for neck, upper back, lower back, chest and abdomen. Note that the pelvic region is greyed out to indicate it was not part of this collection.

**Table 2.** Demographics and pain variables by group (average±standard deviation). Variables are labeled as baseline, 6mo (6-months post-surgery), or 6mo-basline (6 months minus baseline).

| Variable | #<br>NI | # I | Mean (SD) –<br>NI | Mean (SD) – I | p |
| --- | --- | --- | --- | --- | --- |
| Age | 27 | 27 | 40.59 (4.97) | 40.74 (4.92) | 0.917 |
| Pain intensity – baseline | 27 | 27 | 5.7 (1.68) | 5.67 (2.65) | 0.732 |
| Pain intensity – 6 months | 27 | 27 | 0.81 (1.14) | 0.11 (0.58) | < .001 |
| Change in pain intensity<br>(6 mo – baseline) | 27 | 27 | -4.89 (1.63) | -5.56 (2.56) | 0.404 |
| FM score (baseline) | 27 | 27 | 9.67 (4.31) | 9.37 (4.02) | 0.896 |
| FM score (6mo) | 27 | 27 | 8.74 (4.36) | 2.7 (1.79) | < .001 |
| ΔFM score (6mo-baseline) | 27 | 27 | -0.93 (3.68) | -6.67 (3.54) | < .001 |
| Widespread Pain Index (baseline) | 27 | 27 | 2.48 (2.23) | 2.59 (2.42) | 0.979 |
| Widespread Pain Index (6mo) | 26 | 24 | 2.38 (3.25) | 0.42 (0.72) | 0.004 |
| $\Delta$ Widespread Pain Index<br>(6mo – baseline) | 26 | 24 | 0.08 (3.26) | -2.04 (2.31) | 0.044 |
| Pain interference (baseline) | 27 | 27 | 23.48 (8.3) | 21.59 (9.3) | 0.435 |
| Sensory sensitivity (baseline) | 27 | 27 | 1.15 (1.41) | 0.56 (1.15) | 0.044 |

*p-values from Wilcoxon rank-sum tests.*

### SEED-TO-VOXEL FUNCTIONAL CONNECTIVITY

Of the 12 seeds tested (Table 1), no significant group differences were observed during rest. Four seeds showed significant group differences during tonic pressure pain (“pain”), and two seeds showed a significant group difference in the painful pressure-versus-rest contrast (“pain vs rest”) (Table 3).

**Table 3.**
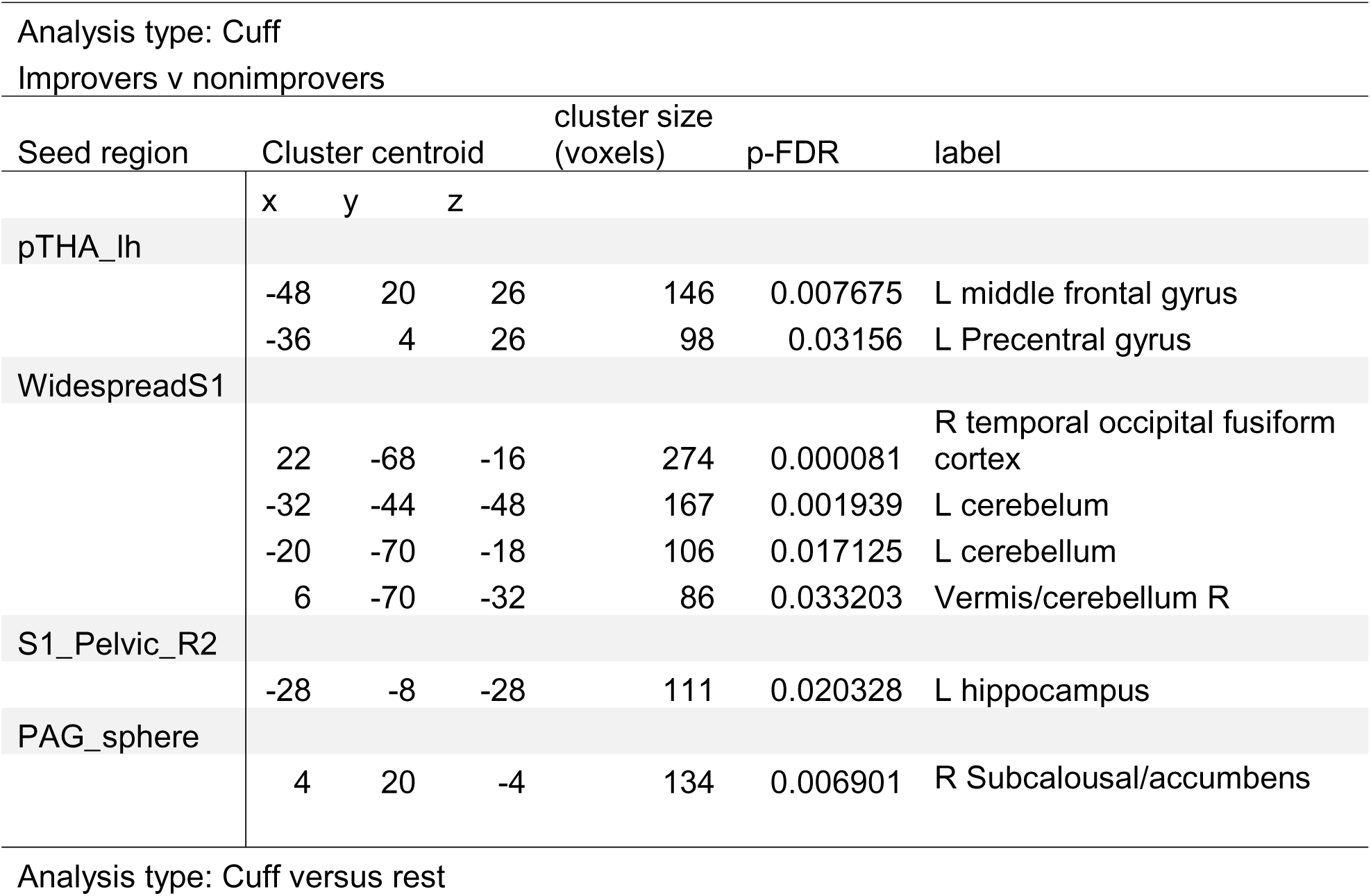
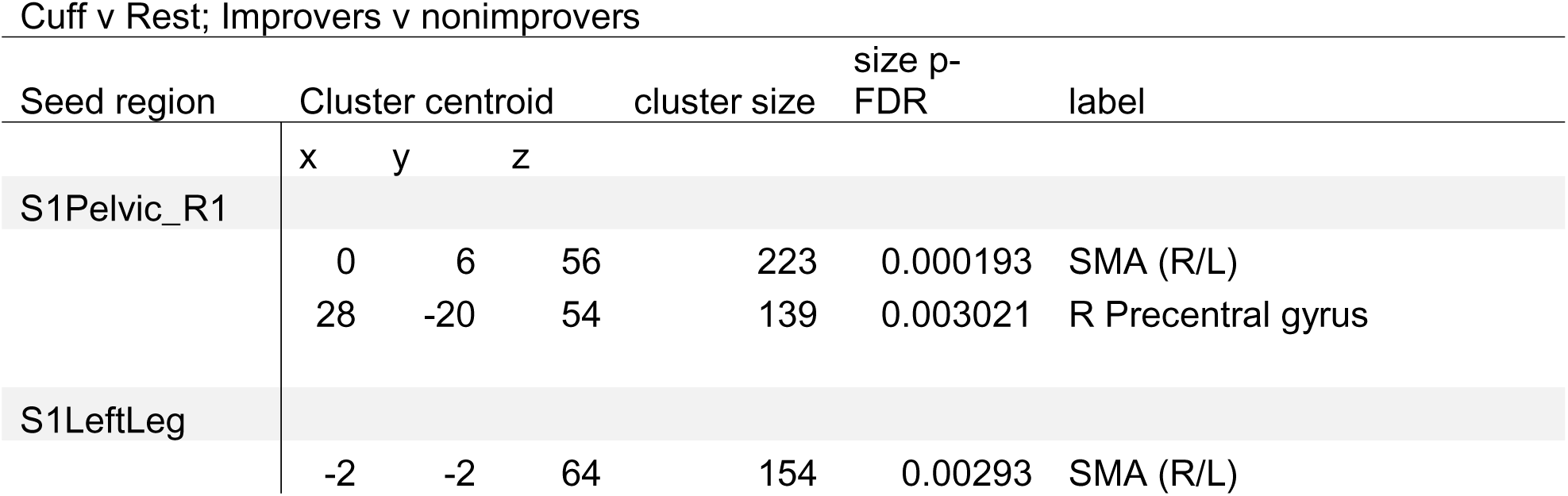
Information for significant clusters identified in the seed-to-whole brain analysis. Clusters are reported separately for the sustained painful pressure analysis and the painful pressure versus rest analysis. Includes cluster centroid, size, and adjusted p-value. Labels were derived from the Harvard-Oxford Cortical and Subcortical atlases and visual inspection for confirmation.

During pain, non-improvers showed greater functional connectivity than improvers between the left posterior thalamus seed (Table 1, seed 7) and clusters in the left middle frontal gyrus and left motor cortex (Figure 2b). Non-improvers also showed greater connectivity between the S1 widespread-pain seed (Table 1, seed 1) and clusters in the left cerebellum, vermis, and right temporo-occipital fusiform cortex (Table 3). A second S1 finding was observed between one of the right pelvic representation seeds (Table 1, seed 3) and the left hippocampus (Figure 2a), with greater connectivity again observed in non-improvers. In contrast, non-improvers showed lower connectivity than improvers between the PAG centroid seed (Table 1, seed 11) and a right subgenual cingulate cluster (Figure 2c).

**Figure 2.**
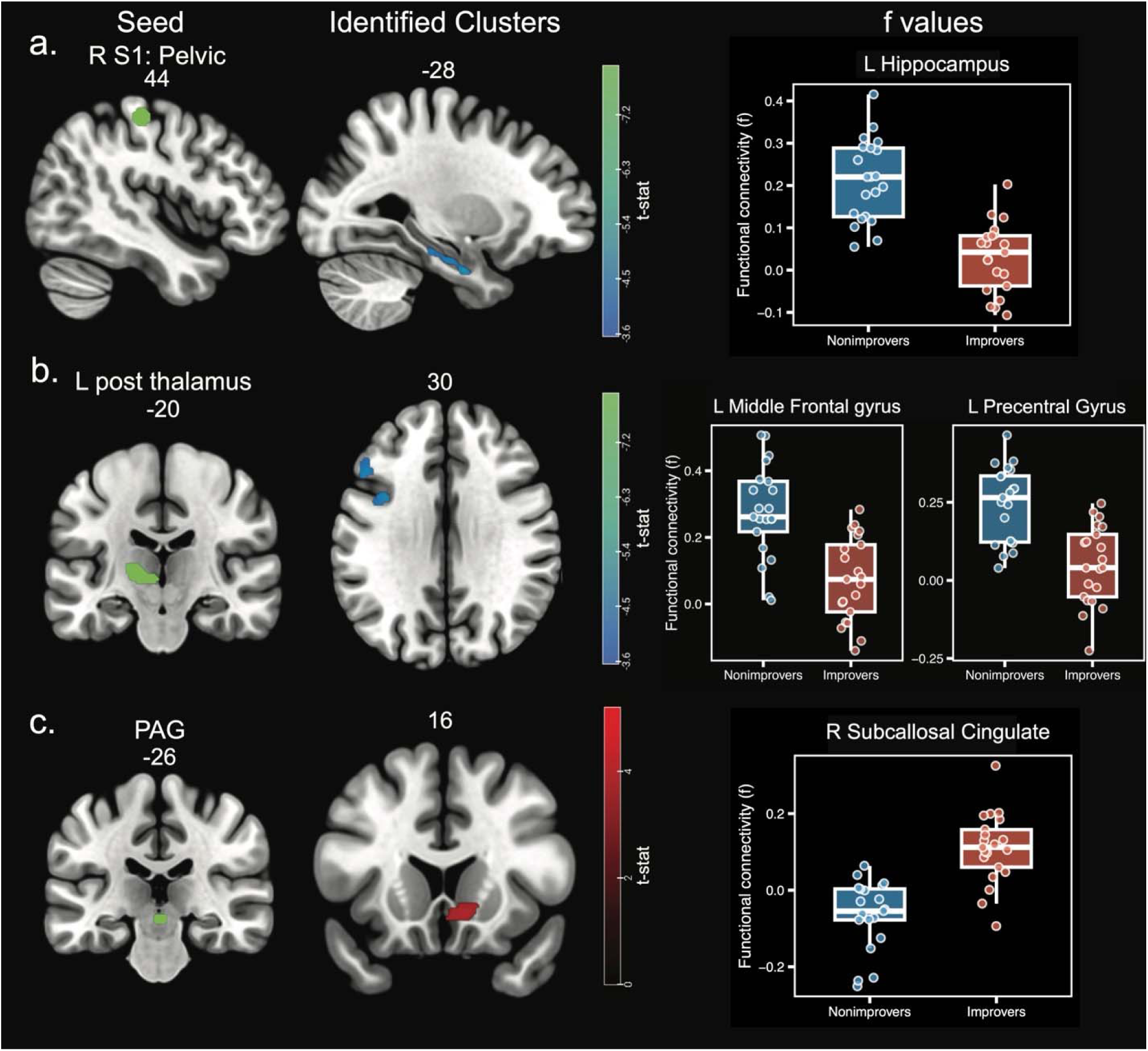
Characterizing improvers and non-improvers. Panel **a** shows individual trajectories of the Fibromyalgia Survey Score (FSS) from baseline to 6 months after hysterectomy for every participant in the improver and non-improver groups. Panel **b** shows the percent of individuals in each group endorsing pain in a regionalized body map before and 6 months following hysterectomy. The map is based on the Widespread Pain Index (WPI), which is made up of 19 sites: Right and left jaw, shoulder, upper arm, lower arm, hip/buttock/trochantor, upper leg, and lower leg, as well as single regions for neck, upper back, lower back, chest and abdomen. Note that the pelvic region is greyed out to indicate it was not part of this collection.

In the painful pressure versus rest contrast, improvers showed a greater decrease in functional connectivity from rest to pressure pain between a right S1 pelvic-representation seed (Table 1, seed 2) and clusters in bilateral supplementary motor area and right motor cortex (Figure 3a). Improvers also showed a greater decrease in functional connectivity from rest to pressure pain between the S1 left leg representation seed (Table 1, seed 5), which corresponds to the cortical representation of the stimulated limb, and clusters in the bilateral supplementary motor area and left lateral occipital cortex (Figure 3b). For both seeds, improvers showed decreased connectivity during pain compared to rest, while non-improvers showed little change or slight increases (Figure 3). There were no clusters identified for the left arm seed.

**Figure 3.**
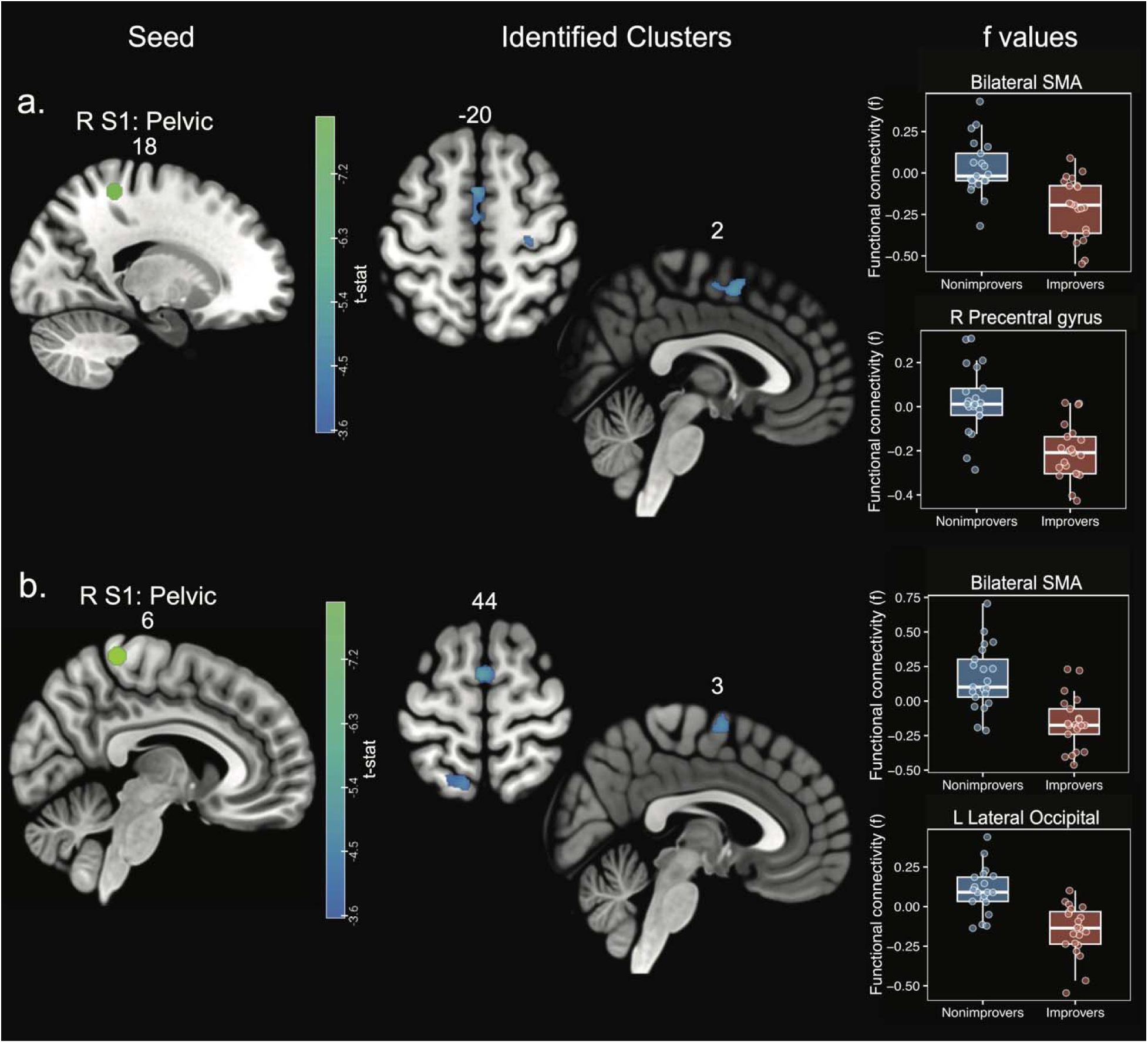
Significant group differences in the change in functional connectivity from rest to sustained painful pressure (painful pressure versus rest analysis). Panels are laid out as in Figure 2. Panel a: a right S1 pelvic representation seed (Table 1, seed 2) and clusters in the bilateral supplementary motor area (SMA) and the right precentral gyrus/motor cortex. Panel b: the S1 left leg representation seed (Table 1, seed 5), corresponding to the cortical representation of the stimulated limb, and clusters in the bilateral SMA and the left lateral occipital cortex. Box-and-whisker plots show the difference in functional connectivity values (painful pressure minus rest) extracted from each cluster for participants in the improver and non-improver groups. Slice coordinates are given in MNI space; full cluster details are reported in Table 3.

### POST HOC CLINICAL ASSOCIATIONS

Functional connectivity values from all significant clusters were extracted and examined in relation to three baseline clinical measures: pain interference, sensory sensitivity, and number of painful body regions. Across these exploratory analyses, five significant relationships were identified in improvers and one was identified in non-improvers (Table 4).

**Table 4.**
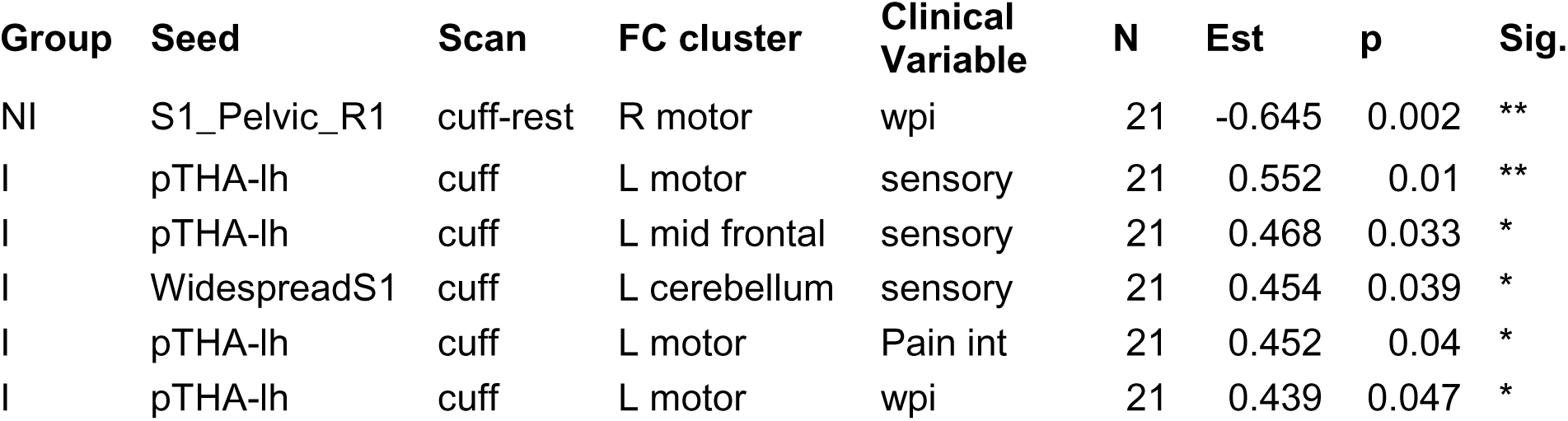
Significant spearman rank correlations between clinical variables at baseline and functional connectivity values extracted from significant clusters. Wpi=widespread pain index; sensory=sensory sensitivity; pain int = pain interference. Seeds are those listed in Table 1. S1_Pelvic_R1= S1 pelvic representation seed in the right hemisphere; pTHA-lh=posterior thalamus, left hemisphere; WidespreadS1=S1 seed identified in previous analyses of widespread pain. Significant values are marked as: * *p* < .05 ** *p* < .01 *** *p* < .001. A full list of tested correlations can be found in Supplementary Table 1.

In the non-improvers, the only significant association was a moderate negative correlation between the number of painful body regions and the pain vs rest connectivity change from the right pelvic S1 (seed 2) to right motor cortex (r = -0.65, p = 0.0016). In the improvers, the strongest association was a moderate positive correlation between left posterior thalamus-to-left motor cortex connectivity during the pressure pain condition and sensory sensitivity scores (*r* = 0.552, *p* = 0.00953). This connection was also significantly associated with pain interference (*r* = 0.452, *p* = 0.0399) and widespread pain (*r* = 0.439, *p* = 0.0466) in improvers. Other significant findings in the improvers group: posterior thalamus-left middle frontal connectivity was associated with greater sensory sensitivity (*r* = 0.468, *p* = 0.0325) and S1-left cerebellar connectivity correlated positively with sensory sensitivity (*r* = 0.454, *p* = 0.0388). All significant brain-clinical associations are reported in Table 4.

## Discussion

Here, we demonstrate that individuals with CPP who subsequently do not improve in nociplastic symptom burden following hysterectomy for benign indications differ presurgically in pain-evoked connectivity. Individuals with CPP who did not see improvement in nociplastic pain symptoms after hysterectomy had differences in ascending sensory and descending modulatory circuitry at baseline: greater pain-evoked recruitment of posterior thalamic and S1-centered circuitry, alongside weaker PAG-subgenual coupling. Overall, the neuroimaging findings suggest that in non-improvers (<50% reduction in FSC), pelvic nociceptive input more readily engages a distributed sensorimotor-surveillance architecture than improvers. These findings align well with the conceptualization of top-down and bottom-up subtypes in nociplastic pain conditions.^8-10^

Nociplastic pain is characterized by hallmark symptoms that extend beyond pain such as spread of pain beyond the primary site, cognitive interference, and issues with sleep.^8^ In a bottom-up presentation, however, peripheral nociceptive drive appears to sustain central sensitization, such that removing the nociceptive source may improve both local and more widespread symptoms such as the ones mentioned above. In a top-down presentation, symptoms appear more centrally maintained and less likely to normalize after a peripheral intervention.^9,11,44^ In osteoarthritis, for example, improvement in nociplastic-associated symptoms after arthroplasty is common, but far from universal, suggesting meaningful neurobiological heterogeneity in surgery responsiveness.^9^ In the present study, individuals were matched on their presurgical FSC scores, and therefore differences in outcome following surgery cannot be explained by preexisting propensity for nociplastic pain symptoms. Indeed, preoperatively, improvers and non-improvers were nearly identical in FSC score, age, the number of painful body sites, and pain interference (Table 2). By design, half of our sample saw greater than 50% reduction in their FSC score following surgery, suggesting their nociplastic symptoms were driven by the purported source of peripheral nociceptive input (i.e. the uterus). This study extends the existing framework of top-down and bottom-up subtypes through the addition of imaging data, allowing us to measure differences in brain function before individuals undergo surgery and experience differences in their nociplastic symptom resolution.

During painful pressure stimulation, non-improvers saw increased functional connectivity between an S1 pelvic representation and the hippocampus (Figure 2a). This may indicate strong neural architecture tying painful stimulation more generally into the pain history associated with CPP. Non-improvers also displayed increased connectivity between the thalamus and left motor cortex (Figure 2b). Interestingly, the motor cortex cluster aligns with one of the recently proposed inter-effector regions of the motor homunculus, which may be involved with whole-body action planning and the somatocognitive action network.^45^ In that context, stronger posterior thalamus-to-precentral coupling in non-improvers is consistent with nociceptive input being routed more strongly into action-oriented and physiologic control circuitry. This may be one way in which pain is maintained as a generalized “body problem” rather than a circumscribed pelvic nociceptive signal. This further aligns with the lack of improvement in whole-body pain in non-improvers (Figure 1, Table 2) and the post-hoc association of these functional connectivity values with widespread pain in non-improvers (Table 4).

The PAG finding is the clearest candidate marker of a stronger, adaptive modulatory system in improvers and aligns with existing literature in CPP. During painful pressure, improvers showed greater connectivity than non-improvers between the PAG centroid seed and a right subgenual cingulate cluster (Figure 2c). The PAG is a central hub of descending pain modulation, and connectivity between PAG and the subgenual cingulate is directly involved with descending pain inhibition.^41-44^ Thus, stronger PAG-subgenual coupling in improvers is consistent with more intact recruitment of descending modulatory and pain-relief circuitry when nociceptive input is encountered. This directly aligns with previous evidence from a voxel-based morphometry study of women with and without endometriosis-associated CPP.^50^ Individuals who had surgically confirmed endometriosis but reported little or no pelvic pain showed increased PAG grey matter volume relative to age-matched controls, whereas women with CPP, with or without endometriosis, instead showed grey matter decreases in the thalamus and other pain-processing regions. Further, in the pain-free endometriosis group, PAG volume trended toward a positive association with the pressure required to elicit mild pain. This could be a sign of heightened antinociceptive activity that allows some women to tolerate an ongoing peripheral nociceptive source without developing chronic pain. Our result extends that hypothesis from structure to function, and from current pain status to treatment outcome: improvers who had stronger PAG-subgenual coupling resemble the adaptive phenotype described above, whereas non-improvers show the weaker coupling expected when descending modulation is less able to compensate once the peripheral source is removed.

When comparing pain to rest, the improvers showed reduced connectivity between an S1 pelvic representation, SMA, and the motor cortex (Figure 3a), while non-improvers saw little change. The same pattern was present for the S1 representation of the stimulated limb (left leg), which showed reduced pain-versus-rest connectivity with the SMA in improvers and little change in non-improvers (Figure 3b). By contrast, no significant clusters were identified with the same analysis using an arm representation in S1. Together, this may indicate a difference in response to painful stimuli including but not limited to the chronic pain relevant site (i.e. pelvis). The lack of findings from the arm representation supports this as a true finding and not a S1-wide phenomenon or noise source. That interpretation is consistent with prior work showing abnormal state-dependent modulation of somatosensory connectivity in fibromyalgia, including failure to show the same pain-related reductions in S1 coupling seen in pain-free controls.^12^

Interestingly, no significant group differences in functional connectivity emerged during rest; group differences were only observed during pain and in the pain vs rest contrast. This is perhaps unsurprising, as the top-down versus bottom-up distinction primarily revolves around how the nervous system handles nociceptive input/whether the source of sensitization is peripheral. Individually calibrating the cuff pressure to elicit the same level of pain in each subject fixes the input at a matched perceived intensity, so residual group differences are more attributable to how a common nociceptive input is processed. This is consistent with broader evidence that stimulus- or task-constrained brain states amplify trait-relevant individual differences and reveal brain-behavior relationships that resting-state connectivity misses.^51^9/26/2026 12:38:00 PM Matching our subjects also removed some of the symptom-burden variance that can drive resting-state group differences elsewhere.^12^ Further investigation in larger samples is required to better understand these differences.

The exploratory correlations between baseline functional connectivity and baseline clinical measures suggest that similar circuits may carry different clinical meanings in improvers and non-improvers. In improvers, stronger pain-state connectivity between the posterior thalamus and left precentral gyrus was associated with greater sensory sensitivity and pain interference; posterior thalamus-left middle frontal connectivity was also associated with greater sensory sensitivity; and widespread-pain S1–left cerebellar connectivity correlated positively with sensory sensitivity. These relationships are consistent with the group-level imaging findings: non-improvers as a group had higher connectivity than improvers in these circuits, and within the improvers group, higher connectivity was associated with greater baseline symptoms. One interpretation is that, in improvers, these connections indexed a reversible sensitization state: greater recruitment of thalamocortical, sensorimotor, and cerebellar circuitry during pain reflected more severe baseline symptoms, but this symptom burden remained responsive to removal of pelvic nociceptive input.

Although this study cannot determine clinical application, the results suggest that preoperative identification of patients with a more top-down nociplastic profile could eventually inform surgical decision-making. Accessible proxies for top-down vs bottom-up nociplastic profiles will have to be developed, but some emerging clinical relationships (i.e. sensory sensitivity) are promising.^52^ In our analysis, the only significant, presurgical difference between improvers and non-improvers at baseline was sensory sensitivity. While the groups had very different outcomes following surgery, their presurgical presentation was highly similar along several of the metrics clinicians may use to determine clinical presentation: pain intensity, pain spread, and pain interference. This finding carried over to the imaging data, where three of the six significant post-hoc associations of clinical variables and functional connectivity were with presurgical sensory sensitivity, potentially indicating its strong link to the neural processes underlying differential nociplastic symptom trajectories following surgery.

Clinicians and patients may need to weigh expected pelvic pain relief against the possibility that broader nociplastic pain symptoms will persist after hysterectomy, but prior work is inconsistent on whether presurgical pelvic pathology is associated with persistent pelvic pain,^53,54^ but these studies have generally not examined comorbid symptoms. Additionally, pathology-based approaches may miss patients whose pain is driven by peripheral factors that are difficult to detect on standard evaluation, such as pelvic floor myofascial dysfunction^55,56^ or neuropathic contributions (e.g., pudendal neuralgia^57^). Therefore, better understanding the neural underpinnings of top-down and bottom-up nociplastic pain subtypes may improve patient selection, counseling, and expectations around hysterectomy for CPP.

Several limitations should be considered. The matching procedure is an important study strength: because baseline nociplastic symptom burden was not different between groups, these findings identify neural processes that are not explained by overall nociplastic severity alone. However, the overall sample size is still modest and consequently the clinical correlations were exploratory and should be interpreted with caution. Future studies should examine these associations in a larger sample and account for variables not examined here, including depression, anxiety, co-existing presence and severity of endometriosis, and residual pathology following surgery (i.e. incompletely excised endometriosis, pelvic adhesions), and concurrent oophorectomy with surgically induced menopause, although these were uncommon in our cohort. Similarly, our ROI approach was hypothesis-driven, focusing on regions selected a priori. Our findings therefore speak specifically to these regions, and future whole-brain analyses may reveal additional connections relevant to the top-down, bottom-up framework. Finally, hysterectomy does not remove all possible peripheral contributors to pain, particularly those unrelated to gynecologic pathology, so lack of FSC score improvement may not reflect a purely central mechanism.

In summary, non-improvers showed greater pain-evoked recruitment of posterior thalamic and S1-centered circuitry with motor, cerebellar, hippocampal, and multisensory association regions, alongside weaker PAG-subgenual coupling. Improvers, by contrast, appeared to retain stronger engagement of circuitry more compatible with adaptive descending modulation and symptom reversibility after peripheral intervention. These findings provide a neurobiological framework for understanding why nociplastic symptoms may improve after removal of a peripheral pain generator in some individuals but persist in others. If replicated prospectively, pain-evoked connectivity patterns could help identify mechanistically distinct recovery profiles and inform interventions that complement surgery by targeting persistent central pain processing.

## Supporting information

AllTables

## Acknowledgements

This work was supported by the National Institutes of Health (NIH) Eunice Kennedy Shriver National Institute of Child Health and Human Development (NICHD) (grant numbers R01HD088712 and R01HD117775). This work was additionally supported by funding from the NIH HEAL Initiative Partnerships to Advance INterdisciplinary (PAIN) Training in Clinical Pain Research (T90DE034663). The authors have no conflicts of interest to declare. The authors thank all of the volunteers who participated in the study.

## References

1. Ahangari A. Prevalence of chronic pelvic pain among women: an updated review. Pain Physician. 2014;17(2):E141–147.

2. As-Sanie S, Ross WT, Till SR. Evaluation and Treatment of Chronic Pelvic Pain. Obstet Gynecol. 2026;147(1):21–43. doi:10.1097/AOG.0000000000006123

3. Lamvu G, Carrillo J, Ouyang C, Rapkin A. Chronic Pelvic Pain in Women: A Review. JAMA. 2021;325(23):2381. doi:10.1001/jama.2021.2631

4. Chronic Pelvic Pain: ACOG Practice Bulletin, Number 218. Obstet Gynecol. 2020;135(3):e98-e109. doi:10.1097/AOG.0000000000003716

5. Misal M, Balk EM, Orlando MS, et al. Predictors of Persistent Pain After Hysterectomy for Chronic Pelvic Pain: A Systematic Review. Obstet Gynecol. 2025;146(5):690–699. doi:10.1097/AOG.0000000000006023

6. As-Sanie S, Till SR, Schrepf AD, et al. Incidence and predictors of persistent pelvic pain following hysterectomy in women with chronic pelvic pain. Am J Obstet Gynecol. 2021;225(5):568.e1-568.e11. doi:10.1016/j.ajog.2021.08.038

7. Orr NL, Huang AJ, Liu YD, et al. Association of Central Sensitization Inventory Scores With Pain Outcomes After Endometriosis Surgery. JAMA Netw Open. 2023;6(2):e230780. doi:10.1001/jamanetworkopen.2023.0780

8. Kaplan CM, Kelleher E, Irani A, Schrepf A, Clauw DJ, Harte SE. Deciphering nociplastic pain: clinical features, risk factors and potential mechanisms. Nat Rev Neurol. 2024;20(6):347–363. doi:10.1038/s41582-024-00966-8

9. Schrepf A, Moser S, Harte SE, et al. Top down or bottom up? An observational investigation of improvement in fibromyalgia symptoms following hip and knee replacement. Rheumatology. 2020;59(3):594–602. doi:10.1093/rheumatology/kez303

10. Kutch JJ, Ichesco E, Hampson JP, et al. Brain signature and functional impact of centralized pain: a multidisciplinary approach to the study of chronic pelvic pain (MAPP) network study. Pain. 2017;158(10):1979–1991. doi:10.1097/j.pain.0000000000001001

11. Kaplan CM, Schrepf A, Mawla I, et al. Neurobiological antecedents of multisite pain in children. Pain. 2022;163(4):e596–e603. doi:10.1097/j.pain.0000000000002431

12. Kim J, Loggia ML, Cahalan CM, et al. The Somatosensory Link in Fibromyalgia: Functional Connectivity of the Primary Somatosensory Cortex Is Altered by Sustained Pain and Is Associated With Clinical/Autonomic Dysfunction. Arthritis Rheumatol. 2015;67(5):1395–1405. doi:10.1002/art.39043

13. Lam J, Mårtensson J, Westergren H, Svensson P, Sundgren PC, Alstergren P. Structural MRI findings in the brain related to pain distribution in chronic overlapping pain conditions: An explorative case–control study in females with fibromyalgia, temporomandibular disorder-related chronic pain and pain-free controls. J Oral Rehabil. 2024;51(11):2415–2426. doi:10.1111/joor.13842

14. Qin ZX, Su JJ, He XW, et al. Altered resting-state functional connectivity between subregions in the thalamus and cortex in migraine without aura. Eur J Neurol. 2020;27(11):2233–2241. doi:10.1111/ene.14411

15. Kim DJ, Lim M, Kim JS, Chung CK. Structural and functional thalamocortical connectivity study in female fibromyalgia. Sci Rep. 2021;11(1):23323. doi:10.1038/s41598-021-02616-1

16. Hemington KS, Coulombe MA. The periaqueductal gray and descending pain modulation: why should we study them and what role do they play in chronic pain? J Neurophysiol. 2015;114(4):2080–2083. doi:10.1152/jn.00998.2014

17. Coulombe MA, Lawrence KSt, Moulin DE, et al. Lower Functional Connectivity of the Periaqueductal Gray Is Related to Negative Affect and Clinical Manifestations of Fibromyalgia. Front Neuroanat. 2017;11:47. doi:10.3389/fnana.2017.00047

18. Amtmann D, Cook KF, Jensen MP, et al. Development of a PROMIS item bank to measure pain interference. Pain. 2010;150(1):173–182. doi:10.1016/j.pain.2010.04.025

19. Häuser W, Jung E, Erbslöh-Möller B, et al. Validation of the Fibromyalgia Survey Questionnaire within a Cross-Sectional Survey. Baradaran HR, ed. PLoS ONE. 2012;7(5):e37504. doi:10.1371/journal.pone.0037504

20. Neville SJ, Clauw AD, Moser SE, et al. Association Between the 2011 Fibromyalgia Survey Criteria and Multisite Pain Sensitivity in Knee Osteoarthritis. Clin J Pain. 2018;34(10):909-917. doi:10.1097/AJP.0000000000000619

21. Brummett CM, Janda AM, Schueller CM, et al. Survey Criteria for Fibromyalgia Independently Predict Increased Postoperative Opioid Consumption after Lower-extremity Joint Arthroplasty: A Prospective, Observational Cohort Study. Anesthesiology. 2013;119(6):1434–1443. doi:10.1097/ALN.0b013e3182a8eb1f

22. Janda AM, As-Sanie S, Rajala B, et al. Fibromyalgia Survey Criteria Are Associated with Increased Postoperative Opioid Consumption in Women Undergoing Hysterectomy. Anesthesiology. 2015;122(5):1103–1111. doi:10.1097/ALN.0000000000000637

23. Williams DA, Schilling S. Advances in the Assessment of Fibromyalgia. Rheum Dis Clin N Am. 2009;35(2):339–357. doi:10.1016/j.rdc.2009.05.007

24. Schrepf A, Williams DA, Gallop R, et al. Sensory sensitivity and symptom severity represent unique dimensions of chronic pain: a MAPP Research Network study. Pain. 2018;159(10):2002–2011. doi:10.1097/j.pain.0000000000001299

25. Schrepf A, Hellman KM, Bohnert AM, Williams DA, Tu FF. Generalized sensory sensitivity is associated with comorbid pain symptoms: a replication study in women with dysmenorrhea. Pain. 2023;164(1):142–148. doi:10.1097/j.pain.0000000000002676

26. Wang S, Peterson DJ, Gatenby JC, Li W, Grabowski TJ, Madhyastha TM. Evaluation of Field Map and Nonlinear Registration Methods for Correction of Susceptibility Artifacts in Diffusion MRI. Front Neuroinformatics. 2017;11. doi:10.3389/fninf.2017.00017

27. Huntenburg ulia M. Evaluating Nonlinear Coregistration of BOLD EPI and T1w Images. Master diss. Max Planck Research Group Neuroanatomy and Connectivity, MPI for Human Cognitive and Brain Sciences, Max Planck Society; 2014.

28. Montez DF, Van AN, Miller RL, et al. Using synthetic MR images for distortion correction. Dev Cogn Neurosci. 2023;60:101234. doi:10.1016/j.dcn.2023.101234

29. Whitfield-Gabrieli S, Nieto-Castanon A. *Conn* : A Functional Connectivity Toolbox for Correlated and Anticorrelated Brain Networks. Brain Connect. 2012;2(3):125–141. doi:10.1089/brain.2012.0073

30. Nieto-Castanon A, Whitfield-Gabrieli S. CONN Functional Connectivity Toolbox: RRID SCR_009550, Release 22. 22nd ed. Hilbert Press; 2022. doi:10.56441/hilbertpress.2246.5840

31. Statistical Parametric Mapping. Elsevier; 2007. doi:10.1016/B978-0-12-372560-8.X5000-1

32. Friston KarlJ, Ashburner J, Frith CD, Poline J-B., Heather JD, Frackowiak RSJ. Spatial registration and normalization of images. Hum Brain Mapp. 1995;3(3):165–189. doi:10.1002/hbm.460030303

33. Whitfield-Gabrieli S, Nieto-Castanon A, Ghosh S. Artifact detection tools (ART). Published online 2011.

34. Power JD, Mitra A, Laumann TO, Snyder AZ, Schlaggar BL, Petersen SE. Methods to detect, characterize, and remove motion artifact in resting state fMRI. NeuroImage. 2014;84:320–341. doi:10.1016/j.neuroimage.2013.08.048

35. Nieto-Castanon A. Preparing fMRI Data for Statistical Analysis. arXiv. Preprint posted online 2022. doi:10.48550/ARXIV.2210.13564

36. Friston KJ, Williams S, Howard R, Frackowiak RSJ, Turner R. Movement-Related effects in fMRI time-series. Magn Reson Med. 1996;35(3):346–355. doi:10.1002/mrm.1910350312

37. Hallquist MN, Hwang K, Luna B. The nuisance of nuisance regression: Spectral misspecification in a common approach to resting-state fMRI preprocessing reintroduces noise and obscures functional connectivity. NeuroImage. 2013;82:208–225. doi:10.1016/j.neuroimage.2013.05.116

38. Behzadi Y, Restom K, Liau J, Liu TT. A component based noise correction method (CompCor) for BOLD and perfusion based fMRI. NeuroImage. 2007;37(1):90–101. doi:10.1016/j.neuroimage.2007.04.042

39. Chai XJ, Castañón AN, Öngür D, Whitfield-Gabrieli S. Anticorrelations in resting state networks without global signal regression. NeuroImage. 2012;59(2):1420–1428. doi:10.1016/j.neuroimage.2011.08.048

40. Tian Y, Margulies DS, Breakspear M, Zalesky A. Topographic organization of the human subcortex unveiled with functional connectivity gradients. Nat Neurosci. 2020;23(11):1421–1432. doi:10.1038/s41593-020-00711-6

41. Bagarinao E, Johnson KA, Martucci KT, et al. Preliminary structural MRI based brain classification of chronic pelvic pain: A MAPP network study. Pain. 2014;155(12):2502–2509. doi:10.1016/j.pain.2014.09.002

42. Fenske SJ, Bierer D, Chelimsky G, et al. Sensitivity of functional connectivity to periaqueductal gray localization, with implications for identifying disease-related changes in chronic visceral pain: A MAPP Research Network neuroimaging study. NeuroImage Clin. 2020;28:102443. doi:10.1016/j.nicl.2020.102443

43. Roux FE, Djidjeli I, Durand JB. Functional architecture of the somatosensory homunculus detected by electrostimulation. J Physiol. 2018;596(5):941–956. doi:10.1113/JP275243

44. Harte SE, Harris RE, Clauw DJ. The neurobiology of central sensitization. J Appl Biobehav Res. 2018;23(2):e12137. doi:10.1111/jabr.12137

45. Gordon EM, Chauvin RJ, Van AN, et al. A somato-cognitive action network alternates with effector regions in motor cortex. Nature. 2023;617(7960):351–359. doi:10.1038/s41586-023-05964-2

46. Lançon K, Séguéla P. Dysregulated neuromodulation in the anterior cingulate cortex in chronic pain. Front Pharmacol. 2023;14:1289218. doi:10.3389/fphar.2023.1289218

47. Meeker TJ, Schmid AC, Keaser ML, et al. Tonic pain alters functional connectivity of the descending pain modulatory network involving amygdala, periaqueductal gray, parabrachial nucleus and anterior cingulate cortex. NeuroImage. 2022;256:119278. doi:10.1016/j.neuroimage.2022.119278

48. Van Strien WWJ, Hollmann MW. Pain Perception and Modulation: Fundamental Neurobiology and Recent Advances. Eur J Neurosci. 2025;62(8):e70275. doi:10.1111/ejn.70275

49. Benarroch EE. Involvement of the nucleus accumbens and dopamine system in chronic pain. Neurology. 2016;87(16):1720–1726. doi:10.1212/WNL.0000000000003243

50. As-Sanie S, Harris RE, Napadow V, et al. Changes in regional gray matter volume in women with chronic pelvic pain: A voxel-based morphometry study. Pain. 2012;153(5):1006–1014. doi:10.1016/j.pain.2012.01.032

51. Greene AS, Gao S, Scheinost D, Constable RT. Task-induced brain state manipulation improves prediction of individual traits. Nat Commun. 2018;9(1):2807. doi:10.1038/s41467-018-04920-3

52. Waller N, Harte SE, Harris RE, et al. Visual Hypersensitivity as a Transdiagnostic Marker of Surgical Pain Response in Arthritis and Chronic Pain Syndromes. Arthritis Rheumatol. 2026;78(6):1339–1349. doi:10.1002/art.70042

53. Hillis S, Marchbanks P, Peterson H. The effectiveness of hysterectomy for chronic pelvic pain. Obstet Gynecol. 1995;86(6):941–945. doi:10.1016/0029-7844(95)00304-A

54. Stovall TG, Ling FW, Crawford DA. Hysterectomy for chronic pelvic pain of presumed uterine etiology. Obstet Gynecol. 1990;75(4):676–679.

55. Phan VT, Stratton P, Tandon HK, et al. Widespread myofascial dysfunction and sensitisation in women with endometriosis-associated chronic pelvic pain: A cross-sectional study. Eur J Pain. 2021;25(4):831–840. doi:10.1002/ejp.1713

56. Pastore EA, Katzman WB. Recognizing myofascial pelvic pain in the female patient with chronic pelvic pain. J Obstet Gynecol Neonatal Nurs JOGNN. 2012;41(5):680–691. doi:10.1111/j.1552-6909.2012.01404.x

57. Ünlü Z, Yentur A, Çakil N. Pudendal Nerve Neuropathy: An Unknown-Rare Cause of Pelvic Pain. Arch Rheumatol. 2016;31(1):102–103. doi:10.5606/ArchRheumatol.2016.5727

